# *Vitellogenin* expression in the ovaries of adult honeybee workers provides insights into the evolution of reproductive and social traits

**DOI:** 10.1101/547760

**Authors:** Carlos Antônio Mendes Cardoso-Júnior, Benjamin P. Oldroyd, Isobel Ronai

**Affiliations:** Departamento de Biologia Celular e Bioagentes Patogênicos, Faculdade de Medicina de Ribeirão Preto, Universidade de São Paulo, Ribeirão Preto, Brasil; Behaviour and Genetics of Social Insects Laboratory, Ecology and Evolution, School of Life and Environmental Sciences A12, University of Sydney, Sydney NSW 2006, Australia

**Keywords:** *Apis mellifera*, epigenetic, honey bee, Juvenile hormone, *Vitellogenin receptor (VgR)*, *Krüppel-homolog 1 (Kr-h1)*, *DNA methyltransferase 3 (Dnmt3)* and the *Forkhead box O transcription factor (FoxO)*

## Abstract

Social insects are notable for having two female castes that exhibit extreme differences in their reproductive capacity. The molecular basis of these differences is largely unknown. A protein that likely plays a key role in these differences is Vitellogenin (Vg), a powerful antioxidant and insulin-signalling regulator. Here we investigate how Royal Jelly (the major food of honeybee queens) and queen pheromone (a major regulator of worker fertility), affects the longevity and reproductive status of honey bee workers, the expression of *Vg*, its receptor *VgR* and associated regulatory proteins. We find that *Vg* is expressed in the ovaries of workers and that workers fed a queen diet of Royal Jelly have increased *Vg* expression in the ovaries. Surprisingly, we find that *Vg* expression is not associated with worker ovary activation. Our findings provide further support for the ‘reproductive ground plan hypothesis’ as Vg has acquired non-reproductive functions in honeybee workers.

## Introduction

Eusocial insects provide an interesting exception to the reproduction-longevity trade-off (Blacher *et al*., 2017; Heinze and Schrempf, 2008), for queens are highly fecund yet have extraordinarily long life-spans, whereas workers are typically non-reproductive and have a short lifespan (Winston, 1987). Understanding of the molecular basis and evolutionary origins of the phenotypic divergence between queen and worker castes, despite their development from identical genomes, are major outstanding questions in sociobiology (Ronai *et al*., 2016b).

A likely mechanism by which queens sustain high rates of egg production over their long lifespan is by life-long production of the egg-yolk precursor protein Vitellogenin (Vg) (Corona *et al*., 2007). The Vg protein is synthesised in the fat body and ovaries of most insects (Hagedorn and Kunkel, 1979), and is taken up by the developing oocyte (Raikhel and Dhadialla, 1992). In eusocial insects Vg seems to have acquired additional functions (Corona *et al*., 2013; Nelson *et al*., 2007). Vg acts as an anti-oxidant that helps prevent senescence in queens and workers (Corona *et al*., 2013). In addition, Vg interacts with components of the insulin signalling pathway (Corona *et al*., 2007), which is increasingly recognised as an important regulator of lifespan (van Heemst, 2010) and behavioural maturation (Amdam *et al*., 2004; Nelson *et al*., 2007).

A significant difference between queens and workers is their diet, which likely contributes to their differential longevity. In honeybees, for example, adult queens are fed glandular secretions that contain Royal Jelly (a nutritious food that is rich in carbohydrates, lipids and protein) (Rembold and Dietz, 1966). In contrast, adult workers feed directly on stored honey and pollen, or on regurgitated crop contents provided by other workers (Winston, 1987). Interestingly, if workers are forced to consume small amounts of Royal Jelly in an artificial diet, they develop a more queen-like phenotype, including ovary activation and increased lifespan (Lin and Winston, 1998; Wang *et al*., 2014; Yang *et al*., 2017). The effects of Royal Jelly seem to be associated with the gene networks that regulate reproduction and longevity (Ford, 2013).

Another environmental factor that influences adult honeybee worker reproduction and potentially longevity is their social environment. Worker fertility is influenced by a number of pheromones released by the brood and the queen, one of which is queen mandibular pheromone (QMP). This pheromone signals the queen’s presence to workers and supresses activation of worker ovaries (Hoover *et al*., 2003; Ronai *et al*., 2016a). Furthermore, QMP regulates the expression of reproductive genes, including *Vg* (Fischer and Grozinger, 2008). When workers are exposed to QMP, *Vg* expression is upregulated (Fischer and Grozinger, 2008). High levels of Vg leads to decreased levels of juvenile hormone (JH) in the haemolymph (Guidugli *et al*., 2005a), which delays behavioural maturation and the initiation of foraging (Marco Antonio *et al*., 2008). This suggests that there is an interplay between the levels of JH and Vg within workers, which in turn influence life expectancy (Amdam and Omholt, 2003; Flatt *et al*., 2013).

The Ovarian Ground Plan Hypothesis argues that the gene networks that were once involved in the regulation of reproduction in the ancestors of social insects have been co-opted to facilitate the regulation of behavioural differences between queen and worker castes (Hunt and Amdam, 2005; West-Eberhard, n.d.). *Vg* appears to have been co-opted in this way (Roy-Zokan *et al*., 2015). While in honeybee queens Vg acts as a precursor for the synthesis of egg-yolk, in workers it acts as a precursor for the synthesis of Royal Jelly proteins (Amdam *et al*., 2003) suggesting that the protein’s function has changed from direct maternal investment in eggs to alloparental care (the production of larval food for sisters). More recently the Ovarian Ground Plan Hypothesis has been extended to the Reproductive Ground Plan Hypothesis which argues that genes that once controlled reproduction now influence behavioural phenotypes between worker sub-castes (Amdam *et al*., 2006). Again, Vg appears to have assumed functions that are only indirectly related to reproduction, most notably behavioural maturation (Nelson *et al*., 2007).

Here we determine whether ovary state (activated or non-activated) and expression of *Vg* in the ovaries of adult honeybee workers are affected by environmental factors. We investigate the interaction effect of two environmental factors that have opposite effects on ovary activation: diet (the presence or absence of Royal Jelly (Hoover *et al*., 2006)) and social environment (the presence or absence of QMP (Hoover *et al*., 2003)). We then investigate how these environmental factors affect the expression of *Vg*’s receptor (*VgR*) (Dohanik *et al*., 2018; Guidugli-Lazzarini *et al*., 2008) and three putative regulators of *Vg. A priori, Vg* expression in the ovaries of workers is likely to be influenced by these two environmental factors because its expression is associated with social functions in addition to its canonical reproductive function.

## Material and methods

### Biological material, cage setup and longevity

We obtained age-matched worker honeybees of standard Australian commercial stock (primarily *A. m. ligustica*) by incubating frames of emerging brood from two source colonies overnight at 34.5 °C. Workers from each source colony were randomly allocated to four cages (Rothenbuhler *et al*., 1979) (n = 150 per cage, total of 8 cages) in a three-factor (source colony plus the two treatments) factorial design. Four of the cages contained half a synthetic queen pheromone strip (Bee Boost Pherotech), which delivered approximately 0.5 queen equivalents per day (Pankiw and Page, 2003; Ronai *et al*., 2015), while the other four cages contained no strip (QMP^-^).

Pollen and water were provided *ad libitum*. In addition, four of the cages (2 QMP^+^ and 2 QMP^-^) were fed a Royal Jelly diet of 50% honey:50% Royal Jelly (Royal Jelly Australia, stored frozen), whereas the other four cages (2 QMP^+^ and 2 QMP^-^) were only fed a Control diet of pure honey. Food was replenished when necessary, and the number of dead workers recorded each day (supplementary table 1). Cages were housed in an incubator at 34.5°C in the dark. At seven and fourteen days of age workers were collected and immediately frozen.

In a parallel experiment to assess longevity, two additional source colonies were prepared as above (four cages per source colony). Cages were kept under the same conditions. However, the workers were not collected and instead the number of dead workers was recorded daily until all workers had died.

### Assessment of ovary activation

We dissected and imaged the ovaries from workers under a dissecting microscope (Leica M60). The paired ovaries of the 7-day-old workers were removed and snap frozen. During dissection the activation state of the ovaries was recorded on a three-point scale: 1 - deactivated (thin ovarioles, no visible oocytes and Vg not required by the oocytes), 2 - semi-activated (ridged ovarioles, with oocytes at the vitellarium stage of development and Vg not yet required for oocyte development) and 3 - activated (mature, white oocytes and Vg required for normal oocyte development) (Ronai *et al*., 2016a; Seehuus *et al*., 2007). Our scale is based on the reproductive control points that occur during early and mid-oogenesis in the honeybee worker (Ronai *et al*., 2016b, 2015).

### Expression analysis

The ovaries of the 7-day-old workers were macerated in TRIzol (Invitrogen) and total RNA extracted with the Direct-zol™ RNA Miniprep kit (Zymo Research) according to the manufacturer’s instructions. Each sample consisted of four pairs of non-activated ovaries (deactivated or semi-activated ovaries), these were pooled to obtain a sufficient amount of RNA. Activated ovaries were not combined as there were a limited number and individual ovaries yielded sufficient RNA. The samples were digested with the Turbo DNase Kit (Thermo Fisher Scientific) to remove DNA. RNA concentration was then quantified using a Qubit 2.0 Fluorometer system (Invitrogen). Each RNA sample was diluted to a final concentration of 15 ng/μL with ultrapure water. We used 142.5 ng of each sample to synthesise cDNA using the SuperScript™ III Reverse Transcriptase Kit (Invitrogen) with Oligo (dT) primer. The cDNA was diluted to 2 ng/μL in ultrapure water.

To detect the expression of *Vg* and its receptor *VgR* in the ovaries of workers (see supplementary table 2 for primer sequences and sources) we first performed conventional PCR. To semi-independently validate *Vg* expression we utilised two sets of *Vg* primers, *Vg*P1 (Lourenco *et al*., 2012) and *Vg*P2 (Koywiwattrakul *et al*., 2005), which target different exons within *Vg* (*Vg*P1: exon 3; *Vg*P2: spans exon 5 and 6). PCR reactions were carried out on three different cDNA samples of Royal Jelly fed workers. The cDNA (2 ng) was mixed with 0.2 μL TaqTi enzyme (Fisher Biotec Australia), 3.2 nmol dNTPs, 1.25 pmol of each primer, buffer 1X, 50 nmol MgCl_2_ in a final volume of 20 μL. The following genes were used as positive controls: ribosomal protein 49 (*RP49*, also known as *rpl32*) and elongation factor 1 alpha (*Ef1α*). The negative control gene was yellow fluorescent protein (*YFP*), which is not present in honeybees. Primer sequences and sources are listed in supplementary table 2. The amplification cycles were: 95 °C for 2 min; 40 cycles of 95°C for 15 sec; 60 °C for 15 sec; and 72 °C for 30 sec, 72°C for 5 min. RT-PCR products were ran on a 1% agarose gel and imaged using a Bio-Rad ChemiDoc Touch Imaging System.

We quantified the expression of *Vg* and *VgR* in the ovaries of adult workers with RT-qPCR. In addition, we quantified the expression of three genes that are potential regulators of *Vg* expression in the honeybee: *Krüppel-homolog 1* (*Kr-h1*) (Amdam and Omholt, 2002; Grozinger *et al*., 2003; Jindra *et al*., 2015), *DNA methyltransferase 3* (*Dnmt3*) (Cardoso-Júnior *et al*., 2018) and *Forkhead box O transcription factor* (*FoxO*) (Corona *et al*., 2007; Hansen *et al*., 2007; Sheng *et al*., 2011; Süren-Castillo *et al*., 2012). RT-qPCR reactions consisted of 2.5 μL SsoAdvanced™ Universal SYBR^®^ Green Supermix (Bio Rad), 1.25 pmol of each primer, 1 μL diluted cDNA (2 ng) in a total volume of 5 μL, with three technical replicates per biological sample. The reactions were performed using a CFX384 Real-Time System (Bio-Rad) with the cycle conditions: 95 °C for 10 min; 40 cycles of 95°C for 10 sec; 60 °C for 10 sec; and 72 °C for 15 sec. For each gene the melt curve was checked to confirm a single amplification peak. The primer efficiency (supplementary table 2) was calculated based on an amplification curve of 10 points obtained through serial dilution of mixed cDNA samples. To normalise the gene expression we used two reference genes (*RP49* and *Ef1α* (Lourenço *et al*., 2008)) whose expression was stable according to *BestKeeper* software (Pfaffl *et al*., 2004). The relative expression was calculated as in (Brito *et al*., 2010) using a relative gene expression formula that takes into account the efficiency of each primer.

### Statistical analysis

Ovary activation score was calculated, per sample, as the average of the activation scores of all ovaries included in the sample. Average ovary activation score and gene expression level were analysed as dependent variables using generalised linear mixed models (GLMM), with ‘colony’ as random effect, and diet and QMP treatments as fixed effects. We used a log function, link = log and family = Gaussian, to model *Vg, VgR* and *Kr-h1* expression irrespective of ovary state. *Dnmt3* and *FoxO* expression and ovary activation score were modelled without applying a log function. Analyses were performed in *R* (Team, 2018) using the packages lme4, car and lsmeans or GraphPad Prism 7. To compare *Vg* gene expression in activated and non-activated ovaries, we used a two-tailed Student’s *t-*test after log^10^ transformation of expression data. Longevity was assessed with survival curves that were statistically compared using Mantel-Cox test. Two-tailed Spearman correlation test was used to assess the correlation between *Vg* and *VgR, Kr-h1, Dnmt3* and *FoxO* expression. A *p*-value lower than 0.05 was considered significant for all statistical tests. When appropriate, significance levels for *post-hoc* tests were Bonferroni adjusted for multiple comparisons.

## Results

### High ovary activation and short lifespan in workers fed Royal Jelly

Ovary activation score was significantly higher in workers fed the Royal Jelly diet compared to workers fed the Control diet (*p* < 0.001; figure 1a and supplementary table 3). In addition, the ovary activation score was significantly higher in workers not exposed to QMP compared to workers exposed (figure 1a, *p* < 0.05, see supplementary table 3 for further details). There was no significant interaction between the diet and QMP treatments (*p* = 0.14; figure 1a). For the majority of the treatment combinations there was a significant difference in ovary activation score when comparing both QMP treatment and diet (comparison of least square means, *p* < 0.001 after Bonferroni correction). The two exceptions were: (i) QMP^+^ and QMP^-^ of the Control diet (*p* = 0.058) and (ii) QMP^-^ of the Control diet and QMP^+^ of Royal Jelly diet (*p* = 0.26). At 14 days of age, workers with activated ovaries comprised 0% of QMP^+^/Control diet, 23% QMP^-^/Control diet, 40% of QMP^+^/Royal Jelly diet and 63% QMP^-^ /Royal Jelly diet (supplementary figure 1).

**Figure 1.**
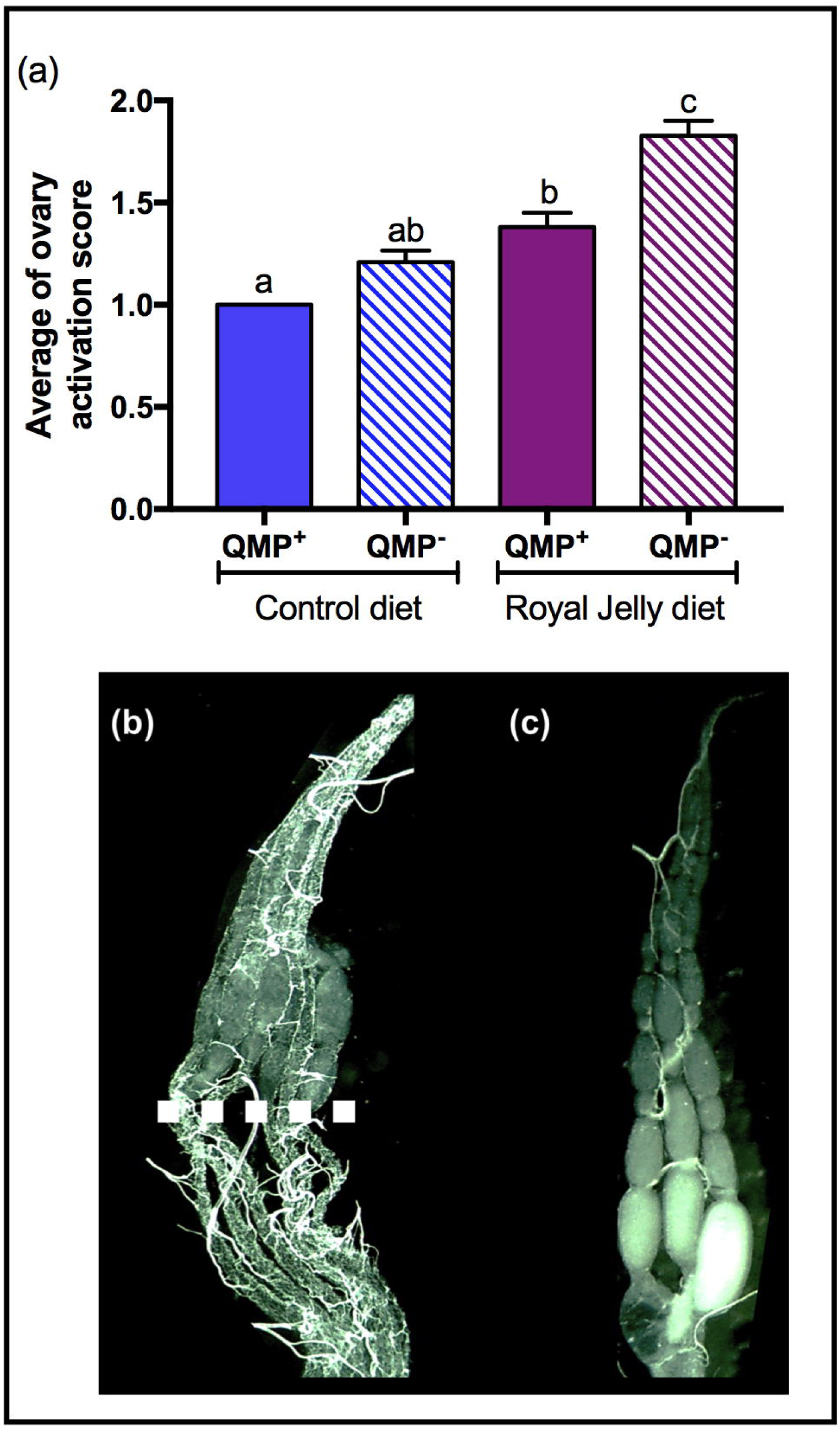
Effect of QMP and Royal Jelly diet on the ovary state of honeybee workers. (*a*) Average ovary activation score from 7-day-old workers treated with QMP and Royal Jelly diet (QMP^+^/Control diet N = 64; QMP^-^/Control diet N = 62; QMP^+^/Royal Jelly diet N = 63; QMP^-^/Royal Jelly diet N = 75; different letters represent *p* < 0.05). See supplementary table 2 for further statistical details. (*b*) Semi-activated ovary from a QMP^+^/Royal Jelly diet worker which shows that Royal Jelly allows the young germ cells to develop whereas the older germ cells have degenerated (dashed line delineates transition). (*c*) Activated ovaries from a QMP^+^/Royal Jelly diet worker.

The Royal Jelly diet reduced the lifespan of workers in both colonies (*p* < 0.0001 Mantel-Cox test followed by Bonferroni correction for multiple comparisons, figure 2). Differences between workers exposed or not to QMP was only found in bees fed Royal Jelly from source colony A (*p* < 0.0001 Mantel-Cox test followed by Bonferroni correction for multiple comparisons, figure 2a).

**Figure 2.**
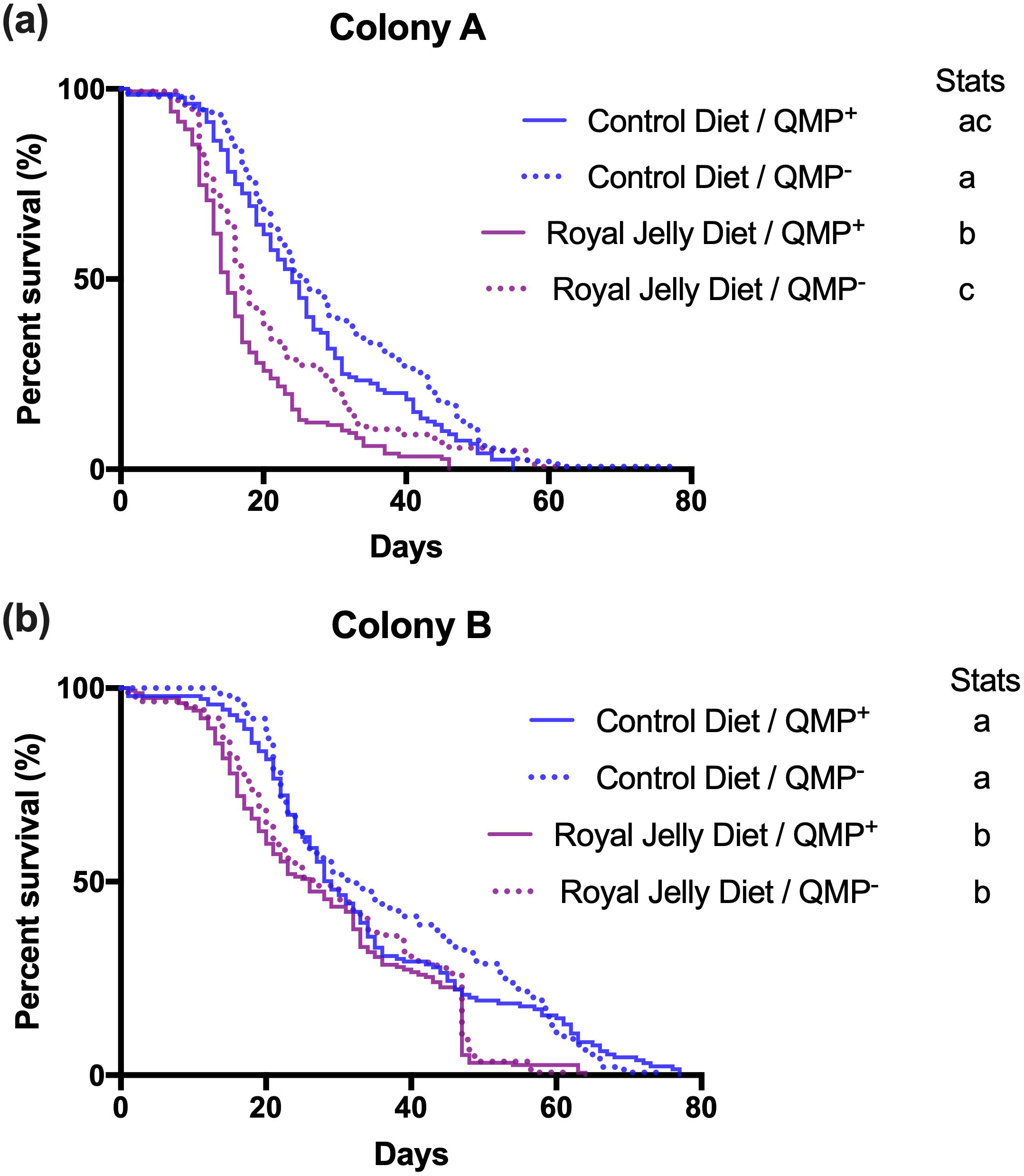
Effect of Royal Jelly and QMP exposure on the longevity of honeybee workers. Longevity rate of workers from two source colonies, Colony A (*a*) and Colony B (*b*) fed Royal Jelly (purple lines) or control diet (blue lines), exposed to QMP (continuous lines) or not exposed to QMP (dashed lines). Different letters represent statistical significance (*p* < 0.05; see the supplementary tables for further details).

### Vg expression in the ovaries of workers is affected by their diet and social environment

Unlike previous studies (Corona *et al*., 2007; Guidugli *et al*., 2005b) we find that *Vg* is expressed in the ovaries of adult honeybee workers (figure 3a and figure 3b). *Vg* expression in the ovaries of adult workers is affected by both QMP (*p* < 0.05; figure 3b and supplementary table 4) and diet (*p* < 0.001; figure 3b and supplementary table 4) with no significant interaction between the two main effects (*p* = 0.8). In the presence of both QMP and the Royal Jelly diet, workers had significantly higher expression of *Vg* in their ovaries compared to the other three treatment groups: QMP^+^/Control diet (comparison of least square mean, *p* < 0.01 after Bonferroni correction); QMP^-^/Control diet (*p* < 0.001) and QMP^-^/Royal Jelly diet (*p* < 0.01). In addition, Vg expression was significantly higher in QMP^-^/Royal Jelly diet workers compared to workers from QMP^-^/Control diet treatment (*p* < 0.01). There was no significant difference in the expression of *Vg* between the two primer sets (*p* = 0.76, *Vg*P1: figure 3b and *Vg*P2: supplementary figure 2).

**Figure 3.**
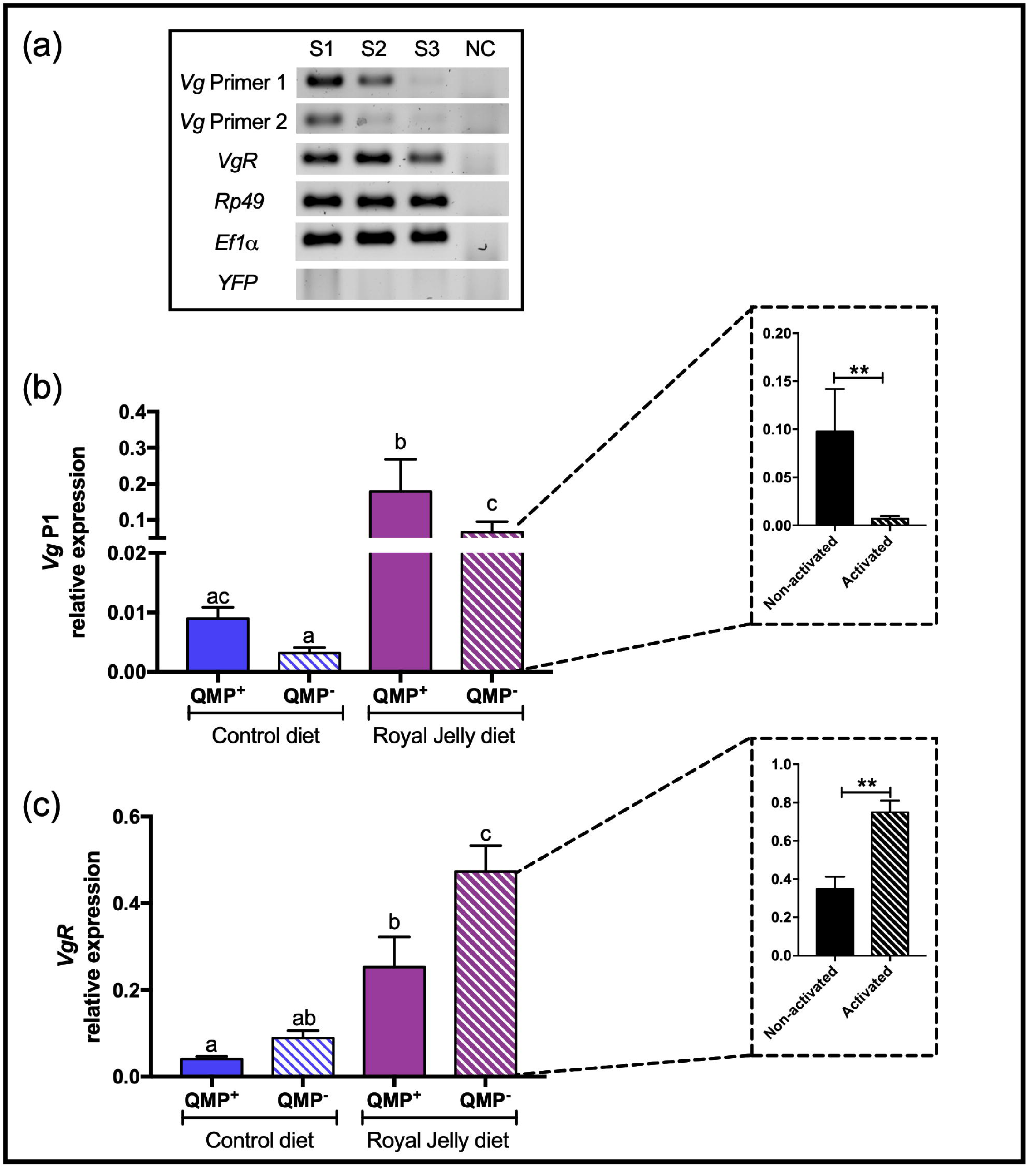
Relative expression of *Vg* and *VgR* in the ovaries of honeybee workers. (*a*) RT-PCR analysis of three samples (S1, S2 and S3) and a negative control (NC) with no cDNA. RP49 and Ef1*μ* were positive control genes and YFP a negative control gene. (*b*) *Vg* (primer set 1) expression in the ovaries of workers exposed to QMP and fed a Royal Jelly or a control diet. Sample size information: QMP^+^/Control diet N = 16; QMP^-^/Control diet N = 15; QMP^+^/Royal Jelly diet N = 18 and QMP^-^/Royal Jelly diet N = 26. The inset box is the QMP^-^ /Royal Jelly diet samples split into the non-activated ovaries and activated ovaries (non-activated ovaries N = 17; activated ovaries N = 9). (*c*) *VgR* expression under the same treatment groups as *Vg*. Different letters represent statistical significance (*p* < 0.05; see the supplementary tables for further details) and ** represents *p* < 0.01 obtained after two-tailed Student’s *t*-tests.

*VgR* expression is affected by both QMP (*p* < 0.05; figure 3c and supplementary table 4) and diet (*p* < 0.001; figure 3c and supplementary table 4) with no significant interaction between the two main effects (*p* = 0.57). *VgR* expression was significantly higher in the treatment combination of Royal Jelly diet with no QMP compared to all other treatment combinations (*p* < 0.01, figure 3c). In addition, *VgR* expression was significantly higher in the treatment with Royal Jelly diet and QMP^+^ compared with the Control diet and QMP^+^ (*p* < 0.001).

### Vg expression is lower in activated ovaries whereas VgR is higher

We investigated the effect of ovary activation on the expression of *Vg* and *VgR* within the QMP^-^/Royal Jelly treatment group to avoid the confounding factors of diet and social environment. *Vg* expression was significantly lower in activated ovaries compared to non-activated ovaries (*p* = 0.004; figure 3b inset, supplementary figure 2 inset and supplementary table 4). In contrast, expression of *VgR* was significantly higher in activated ovaries compared to non-activated ovaries (*p* = 0.0033; figure 3c inset).

### Royal jelly alters the expression of putative Vg regulators

*Vg* expression in the ovaries of workers is not significantly correlated with the expression level of its receptor, *VgR* (*r*^*2*^ = 0.16, *p* = 0.15, supplementary figure 3). To gain insights into the regulatory control of *Vg* transcription in the ovaries of workers, we examined the expression of three additional potential *Vg* regulators. Firstly, the expression of *Kr-h1* was affected by both QMP exposure (*p* < 0.0001, figure 4a and supplementary table 5) and Royal Jelly diet (*p* < 0.0001, figure 4a and supplementary table 5). We found no interaction between the two main effects (*p* = 0.16). Interestingly, the QMP effect was suppressed in the workers fed Royal Jelly (*p* = 0.32). Secondly, the expression of *Dnmt3* is influenced by QMP presence (*p* < 0.0001; figure 4b and supplementary table 5). We found an interaction between QMP and diet (*p* < 0.05). *Dnmt3* is upregulated in the QMP^+^/Control diet in relation to all other groups (comparison of least square mean, *p* < 0.01 after Bonferroni correction). Similar to *Kr-h1*, Royal Jelly suppressed the effects of QMP exposure on *Dnmt3* expression. Thirdly, the expression of *FoxO* was only affected by diet (*p* < 0.0001, figure 4c) with no significant interaction between diet and QMP (*p* = 0.08). *FoxO* expression was higher in QMP^-^/Control diet workers compared to those fed Royal Jelly (comparison of least square mean *p* < 0.05 after Bonferroni correction). In addition, *FoxO* expression was lower in QMP^-^/Royal Jelly diet treatment compared to QMP^+^/Control diet (*p* < 0.0001 after Bonferroni correction).

**Figure 4.**
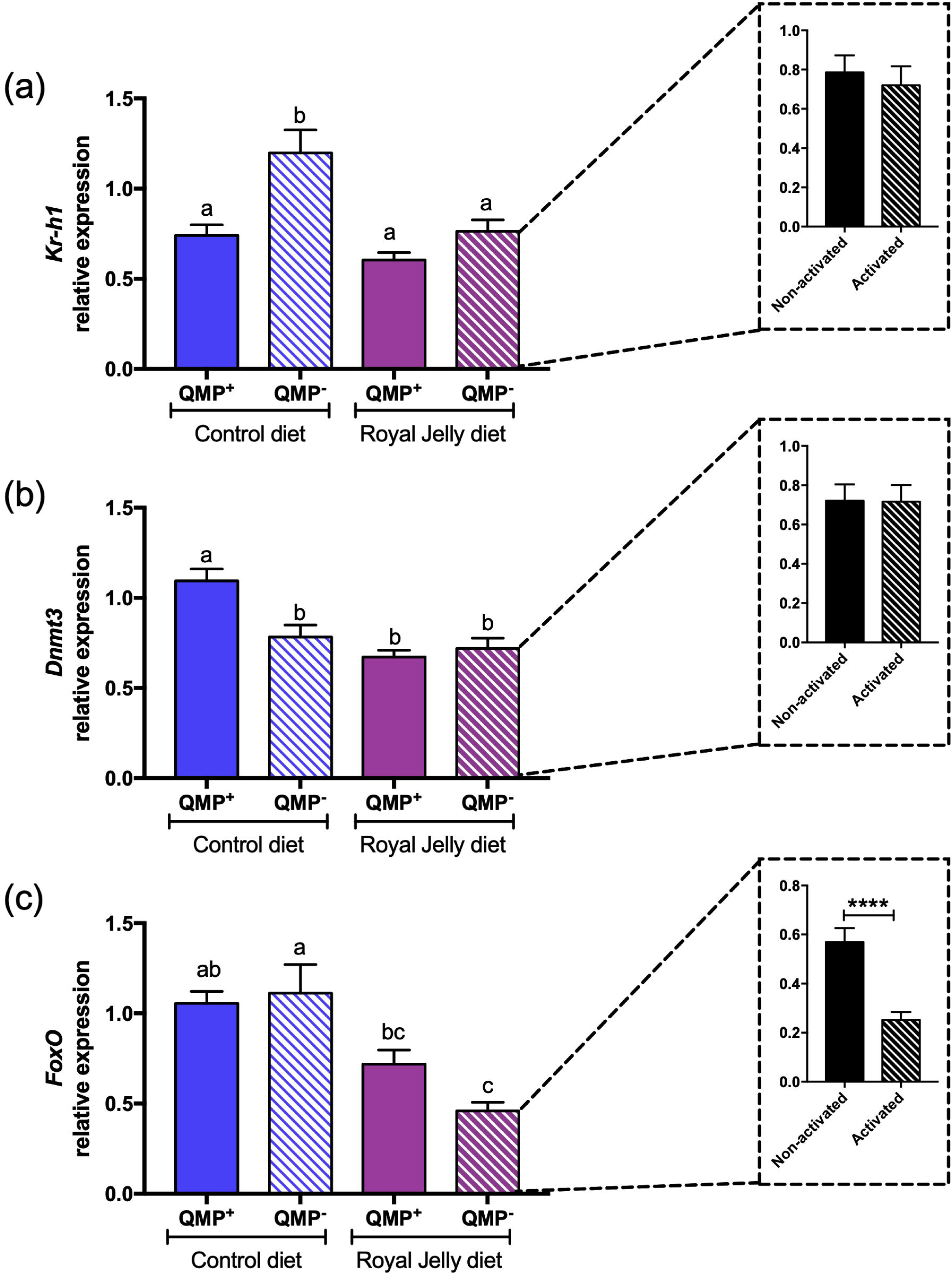
Relative expression of *Kr-h1, Dnmt3* and *FoxO* in the ovaries of honeybee workers. The expression of *Kr-h1* (*a*), *Dnmt3* (*b*) and *FoxO* (*c*) in the ovaries of workers exposed to QMP and fed a Royal Jelly or a control diet. Sample size is the same as in figure 3. Different letters represent statistical significance (*p* < 0.05; see the supplementary tables for further details) and **** represents *p* < 0.0001 obtained after two-tailed Student’s *t*-tests.

We also investigated whether *Vg* expression in the ovaries of workers is correlated with the expression of its potential regulators (supplementary figure 3). *Vg* expression was negatively correlated with the expression of *Kr-h1* (*r*^*2*^ = −0.28, *p* = 0.01, supplementary figure 3) but not correlated with expression of *Dnmt3* (*r*^*2*^ = 0.12, *p* = 0.28, supplementary figure 3) nor expression of *FoxO* (*r*^*2*^ = 0.1, *p* = 0.36, supplementary figure 3).

## Discussion

Our study demonstrates that *Vg*, a gene associated with longevity and behavioural maturation, is expressed in the ovaries of adult honeybee workers. Previously, *Vg* expression was only known in the ovaries of queens, not workers (Corona *et al*., 2007; Guidugli *et al*., 2005b). Honeybee workers are therefore similar to many other social and non-social insects that also express *Vg* in tissues other than the fat body (Jedlicka *et al*., 2016; Valle, 1993). We also find that the environment of a worker (diet and exposure to QMP) influences the expression of *Vg* in its ovaries. While it is known that environmental effects influence *Vg* expression in the fat body of honeybees and other insects (Amsalem *et al*., 2014; Roy-Zokan *et al*., 2015), this is the first report of changes in *Vg* expression in the ovaries.

The decreased expression of *Vg* in activated ovaries compared to non-activated ovaries is unexpected for an egg-yolk precursor protein (Raikhel and Dhadialla, 1992). We propose two plausible explanations. First, in activated ovaries with mature oocytes the Vg protein is translocated from the haemolymph *via* VgR into the oocytes (Dohanik *et al*., 2018; Guidugli-Lazzarini *et al*., 2008). Translocation of large amounts of Vg into the ovaries to nourish developing oocytes may supress transcription of *Vg in situ*. Second, and more speculatively, it is known that Vg influences the population of micro-RNAs in the honeybee brain and fat body, and by this means Vg may influence gene expression in these tissues (Nunes *et al*., 2013). Thus, Vg may be responsible for regulating the microRNA population in ways that control ovary activation state. We therefore suggest that *Vg* has a signalling role in the ovaries of workers, providing additional evidence that the ovary regulates the behaviour of worker honeybees (Corona *et al*., 2007; Page, 2013; Vergoz *et al*., 2012; Wang *et al*., 2010).

Feeding Royal Jelly to workers has a major effect on *Vg* expression in their ovaries (approximately 20-fold higher), which provides an opportunity to investigate the pathways that regulate *Vg* expression in the ovaries. We determined whether genes in physiological, epigenetic and signalling pathways are influenced by the presence of Royal Jelly in the diet. First, *Kr-h1*, a downstream target of JH that is frequently used as a reliable proxy of this hormone (Belles and Santos, 2014; Jindra *et al*., 2015; Minakuchi *et al*., 2009), is downregulated in Royal Jelly-fed workers. This suggests that JH levels are lowered by the presence of Royal Jelly in the diet. Since JH represses *Vg* expression (Amdam and Omholt, 2003), this effect is likely to be a factor that promotes *Vg* transcription in the ovaries of workers fed Royal Jelly. In addition, we find that *Kr-h1* is negatively correlated with *Vg* expression, suggesting that the ovaries play a role in the “double repressor” interaction between JH and Vg that possibly regulates maturation in honeybee workers (Amdam and Omholt, 2003; Cardoso-Júnior *et al*., 2018; Flatt *et al*., 2013). Second, *Dnmt3*, an enzyme involved in DNA methylation, is downregulated in Royal Jelly-fed workers and this diet blocked the effects of QMP. This finding indicates that epigenetic mechanisms may moderate the effects of Royal Jelly in the diet, for both the adult (this study) and larval (Kucharski *et al*., 2008) stages of honeybee workers. Third, *FoxO*, a putative regulator of *Vg* expression in honeybees (Cardoso-Júnior *et al*., 2018; Corona *et al*., 2007), is mainly regulated by post-translational modification and sub-cellular reallocation (van der Heide *et al*., 2004). We found that *FoxO* transcription is reduced in adult workers fed Royal Jelly (this study) as has been previously shown in honeybee larvae (Wheeler *et al*., 2014). We suggest that FoxO negatively regulates *Vg* in adult honeybee workers (but see (Corona *et al*., 2007)). We note that in general, insects fed highly nutritious diets switch on the insulin/IGF1 pathway, which leads to the inactivation of FoxO activity by translocating it to the cytoplasm (Essaghir *et al*., 2009; Sheng *et al*., 2011). Together, these results suggest that Royal Jelly acts through epigenetic, physiological and signalling pathways to increase *Vg* expression.

The diet and social environment of adult honeybee workers have an antagonistic effect on the activation of their ovaries (Lin and Winston, 1998; Ronai *et al*., 2016a). Workers fed a diet containing Royal Jelly activate their ovaries despite the presence of QMP. The Royal Jelly diet appears to counteract the degeneration of the germ cells that normally occurs in the ovaries of workers exposed to QMP (figure 1b, 1c and (Ronai *et al*., 2017, 2015)). Our findings indicate that Royal Jelly can block the inhibitory effect of QMP in the ovaries of workers. As queens have a diet of Royal Jelly, we speculate that queen’s ovaries are protected from her own pheromone by this diet.

The reproduction-longevity trade-off observed in most animals (De Loof, 2011; Hayflick, 2007; Shorter *et al*., 2015) clearly does not apply to honeybees (Winston, 1987). Reproduction may not be costly for queen honeybees since they have unlimited access to a highly nutritious diet of Royal Jelly. If a diet of Royal Jelly increases *Vg* expression in queens, as we have shown here for workers, then high levels of the Vg protein are likely to act as an anti-oxidant (Corona *et al*., 2007; Seehuus *et al*., 2006) and for energy storage (Nilsen *et al*., 2011), key factors in promoting longevity (Remolina and Hughes, 2008). While it is difficult to experimentally test the effects of Royal Jelly in the diet of laying queens, since this is their sole food, our study provides a starting point to uncover the nutrigenomic effects of Royal Jelly in honeybees. Curiously, while a low-dose diet of Royal Jelly increases the lifespan of honeybee workers (Wang *et al*., 2014; Yang *et al*., 2017) and other invertebrates (Honda *et al*., 2011; Wang *et al*., 2015; Xin *et al*., 2016), we found that workers fed a higher dose of Royal Jelly have shorter lifespans. A likely explanation for these contrasting results is that workers cannot tolerate high concentrations of Royal Jelly (Shorter *et al*., 2015; Yang *et al*., 2017). Alternatively, consumption of Royal Jelly may hasten behavioural maturation and workers that are physiologically older are less tolerant of protein in their diets (Paoli *et al*., 2014).

In conclusion, our finding suggests that the role of *Vg* in the non-activated ovaries of adult honeybee workers is not as an egg-yolk precursor protein but for other functions. Our study also provides further support for the ‘reproductive ground plan’ hypothesis, which argues that gene networks that once regulated fertility now regulate behavioural maturation in the worker caste among other functions (Amdam *et al*., 2006; Hunt and Amdam, 2005; West-Eberhard, n.d.).

## Acknowledgments

The authors thank Patsavee Utaipanon for helping with the survival experiment and Melanie Hoang for assisting with ovary dissections. This work was supported by grants from Fundac□ão de Amparo à Pesquisa do Estado de São Paulo (2016/15881-0 and 2017/09269-3 to CA) and the Australian Research Council (DP180101696 to BPO and Amro Zayed).

## Author’s contributions

C.A. designed the study, carried out the dissections, survival experiment and molecular lab work, analysed the data and wrote the manuscript; B.P.O. participated in the design of the study, analysed the data and wrote the manuscript; I.R. conceived the study, designed the study, analysed the data, performed the cage experiment and wrote the manuscript. All authors gave final approval for publication and declare no conflict of interests.

