## Supplementary material for "*Vitellogenin* expression in the ovaries of adult honeybee workers provides insights into the evolution of reproductive and social traits"

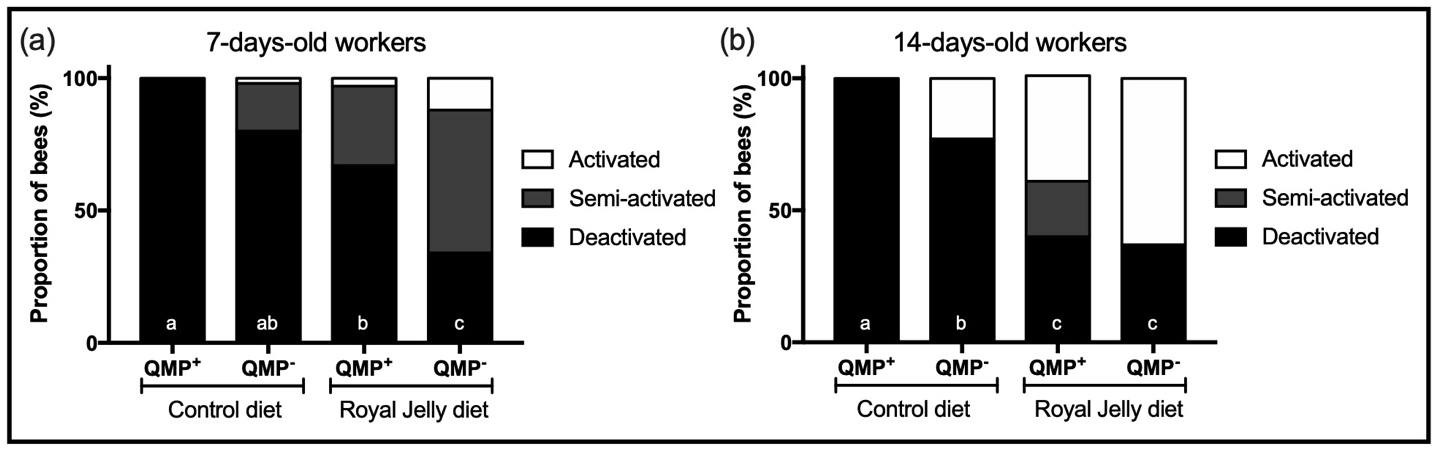

**Supplementary Figure 1.** Effect of QMP and Royal Jelly diet on the ovary state of adult honeybee workers. (*a*) Proportion of 7-days-old workers with deactivated, semi-activated or activated ovaries (QMP^+^/Control diet N = 64; QMP^-^/Control diet N = 62; QMP^+^/Royal Jelly diet N = 63; QMP^-^/Royal Jelly diet N = 75; different letters represent *p* < 0.05). (*b*) Proportion of 14-days-old workers with deactivated, semi-activated or activated ovaries (QMP^+^/Control diet N = 50; QMP^-^/Control diet N = 48; QMP^+^/Royal Jelly diet N = 48; QMP^-^/Royal Jelly diet N = 46; different letters represent *p* < 0.05).

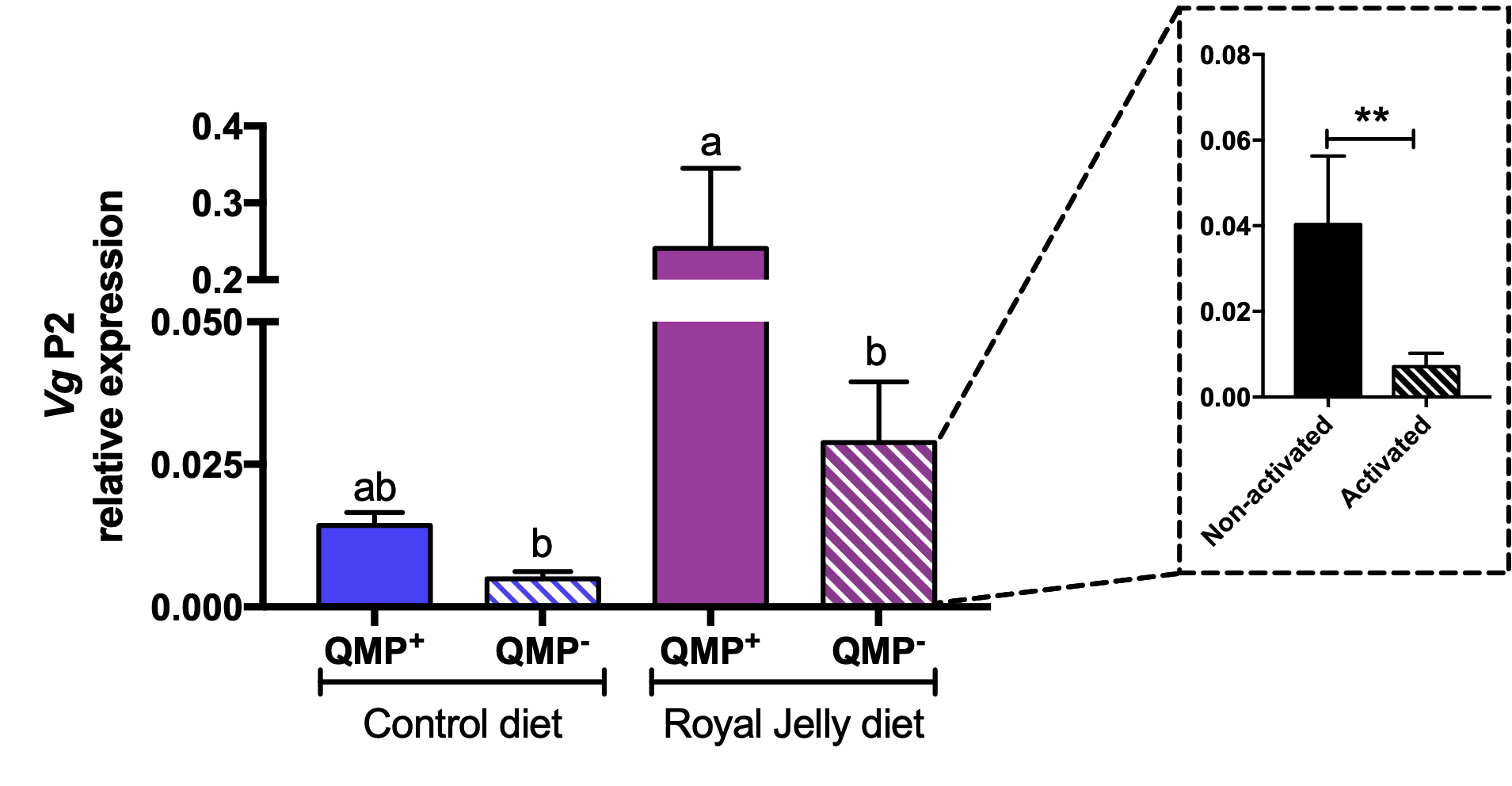

**Supplementary Figure 2.** Expression of *Vg*, using primer set 2, in the ovaries of honeybee workers. Workers were exposed to QMP or not exposed and fed a Royal Jelly or a control diet. Sample size see Figure 3. The inset box is the QMP^-^/Royal Jelly diet samples split into the non-activated ovaries and activated ovaries (non-activated ovaries N = 17; activated ovaries N = 9). Different letters represent statistical significance (*p* < 0.05; see the supplementary tables for further details) and ** represents *p* < 0.01 obtained after two-tailed Student’s *t*-tests.

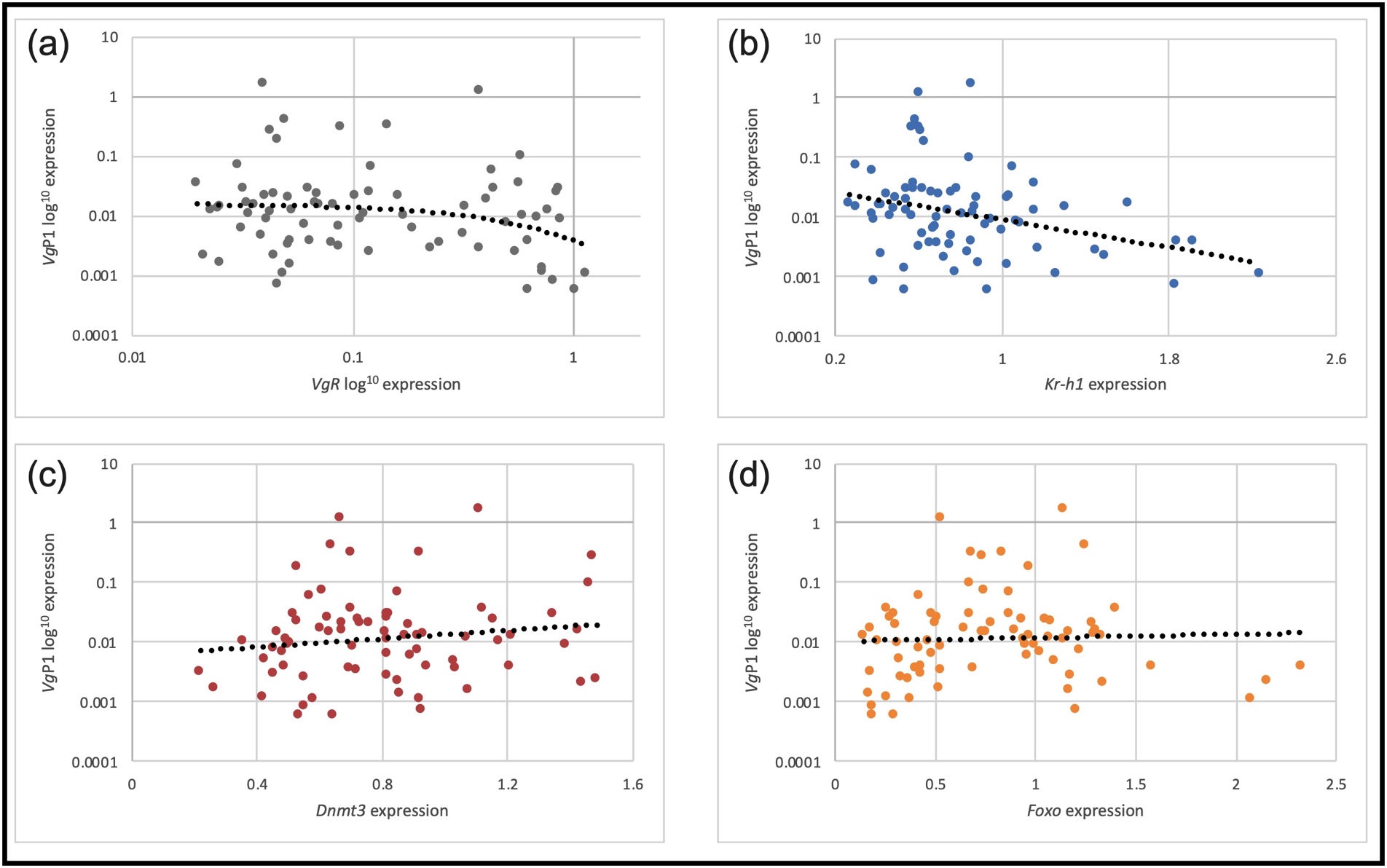

**Supplementary Figure 3.** Correlation analysis between *Vg* and *VgR, Kr-h1, Dnmt3 and FoxO* expression in the ovaries of adult honeybee workers. (a) The expression levels of *Vg* and its receptor, *VgR,* were not correlated (Spearman correlation test R^2^ = -0.1659, *p* = 0.1521, N = 76). (b) The expression levels of *Vg* and *Kr-h1* were negatively correlated (Spearman correlation test R^2^ = -0.289, *p* = 0.0113, N = 76). (c) The expression levels of *Vg* and *Dnmt3* were not correlated (Spearman correlation test R^2^ = 0.1252, *p* = 0.2845, N = 76) (d) The expression levels of *Vg* and *FoxO* were not correlated (Spearman correlation test R^2^ = 0.1044, *p* = 0.3692, N = 76).

**Supplementary table 1.** Number of dead 7-day-old workers in each cage.

| Treatment Group | Replicate cage | Number of dead workers |
| --- | --- | --- |
| QMP^+^/Control diet | 1 | 1 |
|  | 2 | 0 |
| QMP^-^/Control diet | 1 | 5 |
|  | 2 | 3 |
| QMP^+^/Royal Jelly diet | 1 | 1 |
|  | 2 | 0 |
| QMP^-^/Royal Jelly diet | 1 | 8 |
|  | 2 | 4 |

**Supplementary Table 2.** List of primers used in RT-PCR and RT-qPCR. The efficiency of each primer set was used to calculate the relative gene expression; * efficiency used to normalise the expression of *FoxO* and *Kr-h1*.

| Gene | Sequence 5’-3’ | GenBank | Primer efficiency (%) | Reference |
| --- | --- | --- | --- | --- |
| *Vg P1* | F: GCAGAATACATGGACGGTGT | GB49544 | 94 | [1] |
|  | R: GAACAGTCTTCGGAAGCTTG |  |  |  |
| *Vg P2* | F: CCGACGAGGACCTGTTGATTA | GB49544 | 88 | [2] |
|  | R: CTAGGATACGTGGTCATGACA |  |  |  |
| *VgR* | F: ACCTTACGACATTGCCCT | GB40823 | 91 | [1] |
|  | R: TGTGATTTTCGGTCCAAGCCC |  |  |  |
| *RP49* | F: CGTCATATGTTGCCAACTGGT | GB10903 | 74 / 77* | [3] |
|  | R: TTGAGCACGTTCAACAATGG |  |  |  |
| *Ef1α* | F: TGCAACCTACTAAGCCGATG | GB52028 | 80 / 86* | [4] |
|  | R: GACCTTGCCCTGGGTATCTT |  |  |  |
| *YFP* | F: CCTGACAACCACTACCTCAGC | GQ221700 | - | [4] |
|  | R: GAACAGGATCGAGCTGAAGG |  |  |  |
| *Kr-h1* | F: GCACTGGCAGTGACAAGGAA | GB45427 | 76 | [5] |
|  | R: CGTGGAGTGTTATCGTAAGTAGCAA |  |  |  |
| *Dnmt3* | F: CAGCGATGACCTGCGATCGGCGATA | GB55485 | 91 | [6] |
|  | R: TACAGGG TTTATATCGTTCCGAAC |  |  |  |
| *FoxO* | F: TTATGCGAGTGCAGAACGAG | GB48301 | 73 | [7] |
|  | R: AAGCGGACTGTCTGGAAAGA |  |  |  |

**Supplementary Table 3.** Statistical details of GLMM test of ovary scoring data.

| **Test** | **Group** | **Ovary activation score** |
| --- | --- | --- |
| Main effects: GLMM test | QMP | **X^2^ = 6.8035, *p* = 0.009** |
|  | Diet | **X^2^ = 47.901, *p* < 0.0001** |
|  | Interaction QMP*Diet | X^2^ = 2.1289, *p* = 0.14 |
| Least square comparison after Bonferroni correction | QMP^-^/Control x QMP^+^/Control | ***p* = 0.0580** |
|  | QMP^-^/Control x QMP^-^/Royal Jelly | ***p* <0.0001** |
|  | QMP^-^/Control x QMP^+^/Royal Jelly | ***p* = 0.2568** |
|  | QMP^+^/Control x QMP^-^/Royal Jelly | ***p* <0.0001** |
|  | QMP^+^/Control x QMP^+^/Royal Jelly | ***p* <0.0001** |
|  | QMP^-^/Royal Jelly x QMP^+^/Royal Jelly | ***p* <0.0001** |

**Supplementary Table 4.** Statistical details of GLMM test used to model *Vg* P1, *Vg* P2 and *Vg*R.

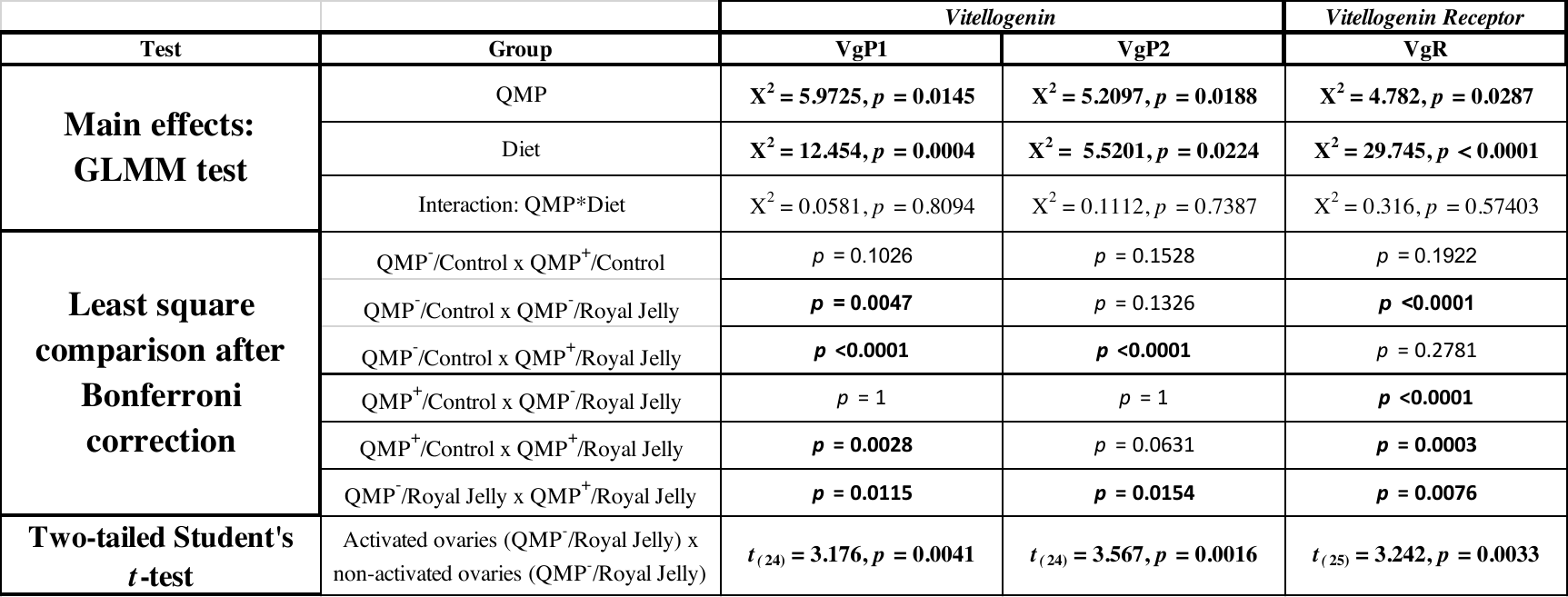

**Supplementary Table 5.** Statistical details of GLMM test used to model *Kr-h1*, *Dnmt3* and *FoxO.*

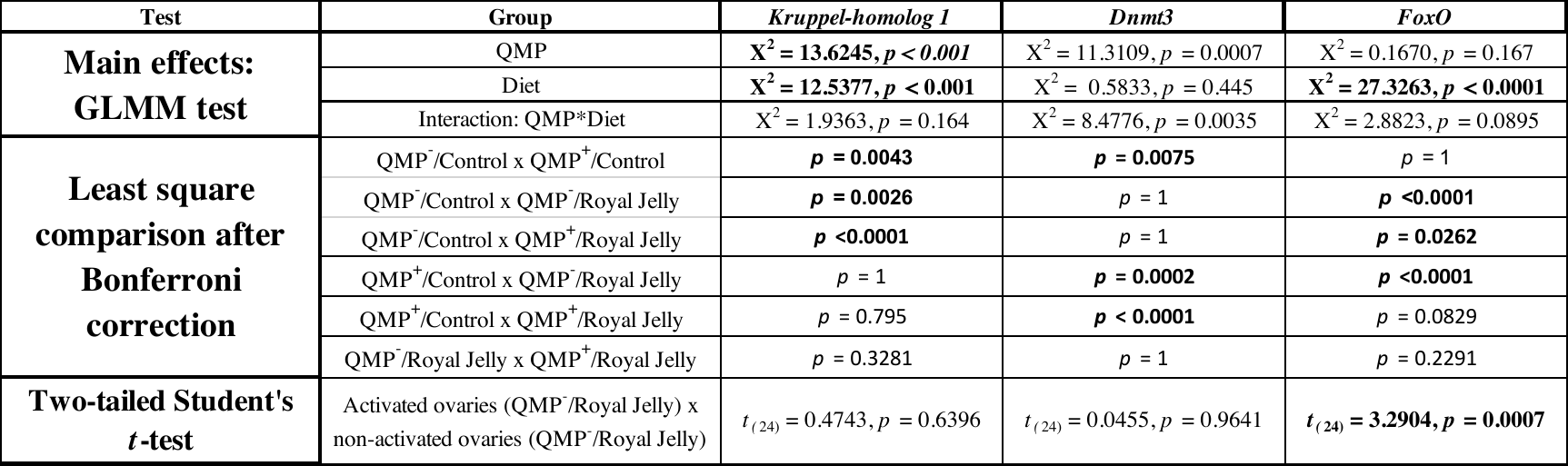
